# A stable reference human transcriptome and proteome as a standard for reproducible omics experiments

**DOI:** 10.1101/2022.11.16.516732

**Authors:** Shaohua Lu, Hong Lu, Tingkai Zheng, Huiming Yuan, Hongli Du, Youhe Gao, Yongtao Liu, Xuanzhen Pan, Wenlu Zhang, Shuying Fu, Zhenghua Sun, Jingjie Jin, Qing-Yu He, Yang Chen, Gong Zhang

**Author notes:** Correspondence: Gong Zhang, E-mail address;, Yang Chen, E-mail address., Shaohua Lu. These authors have contributed equally to this work and share first authorship (Shaohua Lu, Hong Lu, Tingkai Zheng).

## Abstract

In recent years, the development of high-throughput omics technology has greatly promoted the development of biomedicine. However, the poor reproducibility of omics techniques limits its application. It is necessary to use standard reference materials of complex RNAs or proteins to test and calibrate the accuracy and reproducibility of omics workflows. However, the transcriptome and proteome of most cell lines shift during culturing, which limits their applicability to serve as standard samples. In this study, we demonstrated that the human hepatocellular cell line MHCC97H has a very stable transcriptome (R^2^=0.966-0.995) and proteome (R^2^=0.934-0.976 for DDA, R^2^=0.942-0.986 for DIA) after 9 subculturing generations, which allows this stable standard sample to be stably produced on an industrial scale for several decades. Moreover, this stability was maintained across labs and platforms. In sum, our results justified a omics standard reference material and reference datasets for transcriptomic and proteomics research. This helps to further standardize the workflow and data quality of omics techniques and thus promotes the application of omics technology in precision medicine.

## Introduction

The booming applications of omics technologies, including next-generation sequencing and mass spectrometry, provide unprecedented insights of biology and medicine. However, the reproducibility of the omics technology has been questioned since long. In the genomic sequencing field, a study showed zero sensitivity in finding pathogenic mutations using whole exome sequencing of 57 patients (*1*). The mutation of 40 circulating tumor DNA (ctDNA) samples, sequenced by two individual companies, showed only 12% congruence (*2*). In the RNA-seq field, a wide variety of the methodology limits the reproducibility (*3*). In the proteomics field, a study at HUPO tested samples of 20 highly purified recombinant human proteins containing one or more 1250 Da unique pancreatic peptides per protein were distributed to 27 laboratories for identification. Results showed that only 7 of 27 laboratories reported all 20 proteins, whereas only 1 laboratory reported all 1250 Da trypsin peptides (*4*). Another study found that, even using optimal conditions and a uniform SOP, the median value of protein repeatability for the same mixture sample in different laboratories was only 75%. (*5*). In addition, repeatability of multiple quantitative tests on the same sample in the same laboratory and between different laboratories is less than 80% (*6*). Other studies show that the lack of reliability and repeatability of omics research is one of the biggest obstacles to narrowing the gap between personalized medicine research and practice (*7, 8*).

To examine the accuracy and reproducibility of the omics workflow, reference materials are needed. For genome (DNA) sequencing, reference genome samples, e.g. ΦX174 viral DNA and NA12878 human cell line genomic DNA, were widely used as standards. Mixed genomic standards are used for variant calling benchmarks (*9*). DNA standards are easy to produce because of the high fidelity of DNA replication of normal genomes. However, the transcriptome (RNA) and proteome are qualitatively and quantitatively highly variable due to various kinds of factors, including intrinsic factors (e.g. senescence, cell cycles, contact inhibition, etc.) and extrinsic factors (e.g. temperature, osmolarity, buffer content, oxidation, etc.), which set challenges on making reference standard transcriptome and proteome sample. Since 2004, the Universal Human Reference RNA (UHRR) (*10*) has been commercialized as a “standard” RNA sample for RNA-seq benchmarking (*11-15*). However, the RNA content of the UHRR sample may shift during the long-time production. UHRR is a pooled RNA of 10 cancerous cell lines, including HeLa, which is well known for its instability (*16-21*). Indeed, in the lot-to-lot comparison of the UHRR, the Pearson correlation reached only R^2^=0.9478 (R=0.9736, data from (*10*)), which demonstrated such instability even in short production period. Such variation, which is expected to multiply during long production period, is not sufficient to evaluate the reproducibility of the advancing next-generation sequencing techniques with increasing depth and resolution.

The proteome is more variable than the transcriptome due to the massive translational regulation (*22*). Therefore, a stable proteome reference is more difficult to produce. Since 2003, the Proteomics Standards Initiative standardized the data formats of mass spectrometry (MS)-based proteomics, but did not plan to provide a human proteome standard sample (*23, 24*). Till 2021, proteome standard material has not been considered in the quality standards in research facilities (*25*). Currently, a mixture of 18∼48 recombinant proteins are used as a “test standard” or spike-in for proteomics (*5, 26-29*). However, such small number of proteins could hardly form a reference standard for complex proteome samples.

In this study, we exhibited a human cell line with stable transcriptome and proteome across multiple passages, which is suitable as a material for long-term production of reference transcriptome and proteome standard materials.

## Results

### Finding a stable cell line

To maintain the long-term production of reference transcriptome and proteome, a cell line with stable transcriptome and proteome over subculture generations is needed. We tested 5 commonly-used cell lines, MHCC97H, MHCCLM6, MHCCLM3, HeLa and A549. For each cell line, we cultured for 8∼12 generations and took samples from each generation (Fig. 1). Total RNA was extracted from each sample and the RNA quality was examined by electrophoresis to verify that they were not degraded (Supplementary Fig. 1A-E). The polyA+ mRNA was then sequenced and quantified using RPKM method. The mutual correlation among showed that the MHCC97H has the most stable transcriptome, with the Pearson R^2^=0.966∼0.995 (Fig. 2A). The other two hepatocellular carcinoma cell lines showed lower consistency (R^2^ can be as low as 0.947 and 0.921, respectively, Fig. 2B∼C). The HeLa and A549 cell showed even lower consistency over the generations (average R^2^ = 0.955 and 0.923, respectively, with the lowest value 0.899 and 0.846, respectively, Fig. 2D∼F). To be precise, such deviation contains biological deviation (the inconsistency of the transcriptome over generations) and the experimental error (the error generated by the library construction, sequencer and data processing algorithms). We tested also the experimental error by sequencing the same MHCC97H RNA sample 3 times and yielded the Pearson correlation of R^2^=0.955∼0.998 (Fig. 2G). This showed that the deviation shown in Fig. 2A (MHCC97H generations) can be almost explained by the experimental error, i.e. the biological deviation of MHCC97H is almost negligible. In contrast, the lower R^2^ of the other tested cell lines over generations (Fig. 2B∼E) demonstrated their considerable biological deviations over generations, and thus are not ideal candidates for standards.

**Figure 1:**
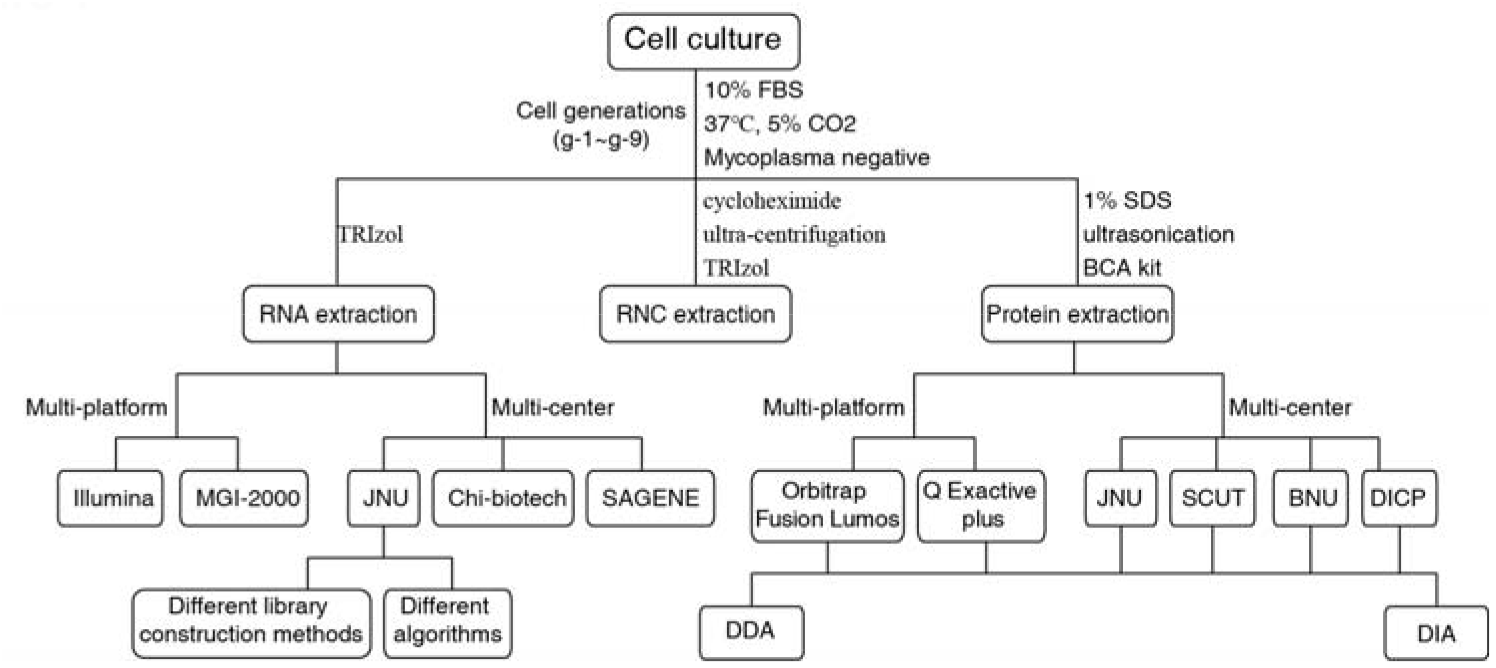
Study design for a stable transcriptome and proteome reference.

**Figure 2:**
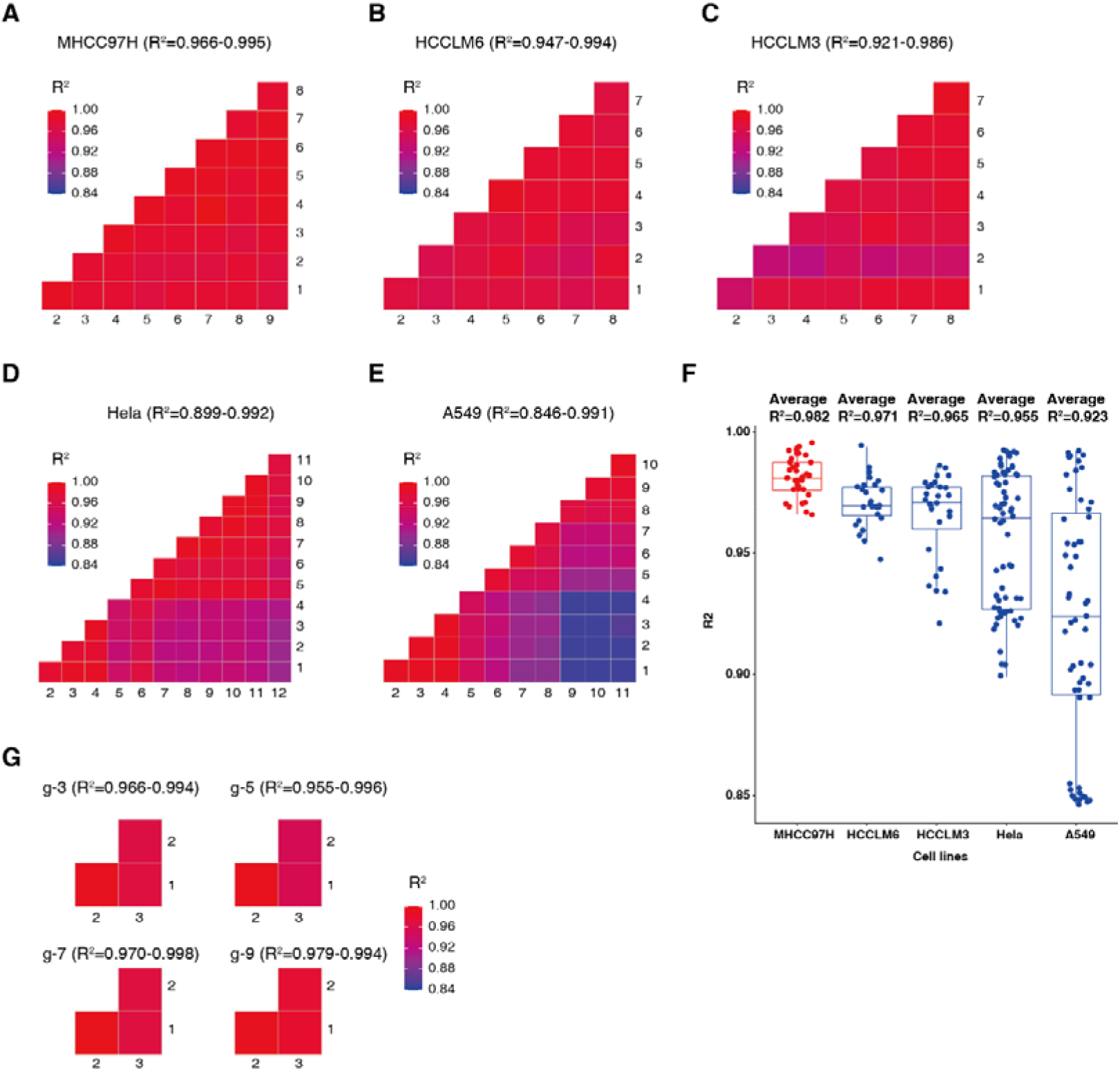
MHCC97H has the strongest transcriptome stability in the subculture of 5 cell lines. (A-E) Mutual Pearson correlation of the transcriptome of MHCC97H, HCCLM6, HCCLM3, Hela, and A549. (F) Distribution of the mutual Pearson correlation of the 5 cell lines. (G) The mutual Pearson correlation of technical triplicates of 4 generations of MHCC97H.

### Robustness of the MHCC97H as a transcriptome reference

A good reference standard should be easy to produce, complemented by a robust standard processing protocol that can yield always the same results, ideally. The major steps introducing deviations are cell culture, mRNA enrichment, library construction/sequencing and data processing algorithms (Fig. 1). First, we subcultured two batches of MHCC97H cell lines in May and December 2021, respectively. The mutual correlation of gene expression over generations was similar (R^2^=0.978 ± 0.013, Fig. 3A). The correlation of the same generation between the two batches are steadily high (R^2^= 0.996 ± 0.002, Fig. 3B). Compared to the technical error (Fig. 2G), the batch variation can be neglected.

**Figure 3:**
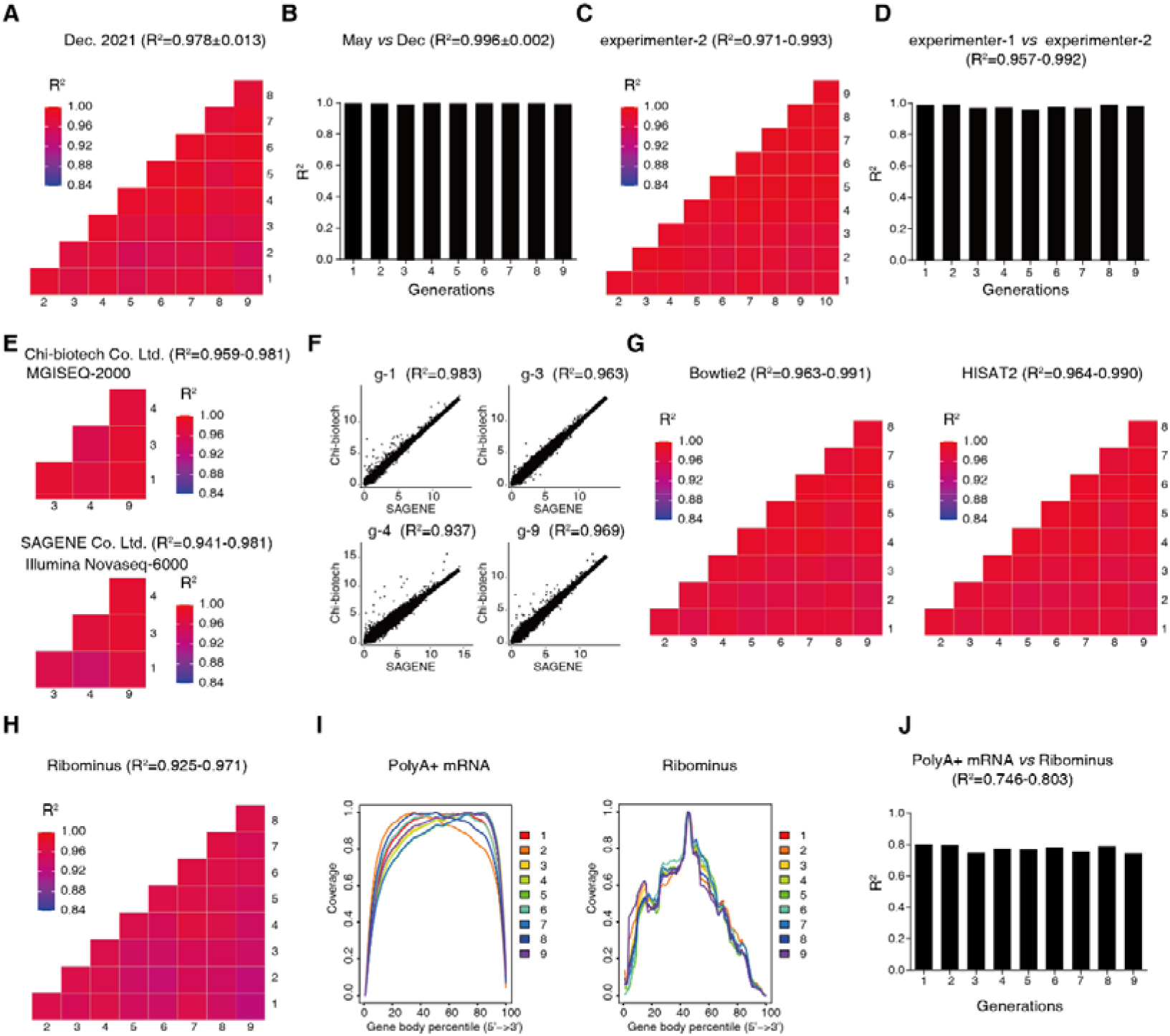
MHCC97H transcriptome was highly stable. (A) The mutual Pearson correlation of the second batch of MHCC97H subcultured in December 2021. The batch shown in Figure 2A was subcultured in May 2021. (B) The correlation of the same generation between the two batches of subcultured MHCC97H cell lines in May and December 2021 (comparing data shown in Figure 2A and Figure 3A). (C) The same series of total MHCC97H RNA was subjected to library construction by another experimenter using another batch of library construction kit. (D) Coincidence of the two experimenter and two batches of library construction kits (comparing data shown in Fig. 3A and Figure 3C). (E) RNA samples from 4 randomly chosen generations (generation 1, 3, 4 and 9) of MHCC97H were distributed to two commercial sequencing service providers. Here shows the mutual Pearson correlation of the transcriptome quantifications within each lab. (F) The cross-lab correlation of gene expression for each generation. (G) Same as Figure 2A. The raw sequencing datasets were analyzed using the widely-used pipelines of Bowtie2 + featureCounts and HISAT2 + featureCounts, respectively. (H) Similar to Figure 2A, using the rRNA depletion strategy (Ribominus) to construct RNA-seq library. (I) Read coverage distribution along the mRNAs in the polyA+ mRNA-seq and Ribominus strategies, respectively. (J) The correlation of gene expression between the PolyA + mRNA and Ribominus strategies for each generation.

Second, we tested the robustness over experimenters and labs. We used the same series of total MHCC97H RNA as starting material, and asked another experimenter to independently constructed the library using another batch of library construction kit. The results (Fig. 3C∼D) were almost identical to the former experimenter (Fig. 2A). We also sent 4 samples to two commercial sequencing service providers more than 1000km away. Chi-Biotech Co. Ltd. was equipped with MGISEQ-2000 sequencer, and Sagene Co. Ltd. was equipped with Illumina NovaSeq-6000 sequencer. They constructed libraries using our protocol and operated the sequencers according to manufacturer’s instructions. The Pearson R^2^ reached 0.959∼0.981 and 0.941∼0.981, respectively (Fig. 3E), and the mutual correlation of gene expression over generations were similar in both labs (Fig. 3F). These results showed that the MHCC97H and our protocol is robust across labs and operators and thus easy to implement.

Next, we investigated the influence of mapping and quantification algorithms on the robustness. Our standard SOP used FANSe3 algorithm and RPKM method to quantify gene expression to reach R^2^=0.966∼0.995 in MHCC97H subculture datasets (Fig. 2A). We also tested two widely-used RNA-seq pipelines, Bowtie2 + featureCounts (*30-35*) and HISAT2 + featureCounts (*36-38*). However, these pipelines showed lower R2 = 0.963∼0.990 (Fig. 3G), showing that these pipelines were less robust than FANSe, which is consistent to the conclusions of previous studies (*39-41*).

We then tested different mRNA enrichment strategies. Our standard protocol used oligo-dT to enrich polyA+ mRNA (mature mRNA), which is applied in most studies. Another strategy is the rRNA depletion strategy (also called “Ribominus”), which removes rRNA by probe hybridization followed by beads extraction or RNaseH degradation. Using rRNA depletion strategy, the Pearson R^2^=0.925∼0.971 (Fig. 3H), which was considerably lower than the polyA+ method. Moreover, used oligo-dT to enrich polyA+ mRNA strategy, the coverage of the reads was much more uniform than the rRNA depletion strategy (Fig. 3I). These showed that the polyA+ method was more robust than the rRNA depletion strategy and should be prioritized when possible. As expected, the correlation of gene expression between the two strategies was not high (R^2^ was only 0.746∼0.803) (Fig. 3J), suggesting that if the data generated by two different strategies should not be mixed for analysis.

### The comparable results of degraded and non-degraded RNA samples

The RNA is vulnerable to the ubiquitous RNases and environmental changes (e.g. freeze-thaw cycles). Therefore, minor or major degradation might be inevitable during the production, storage and transport of the standard samples. We tested the RNA samples under degradation conditions to investigate how the degradation affects their applicability to serve as reference. First, we created a scenario which mimics the degradation due to the environmental exposure: the RNA samples were exposed to the air at room temperature for prolonged time (more than 2 hours), so that the RNases in the environment may enter the tube and degrade the RNA. Then the samples were frozen and thawed for 10 cycles. The electrophoresis showed that the total RNA of MHCC97H was degraded to various extent (Fig. 4A). The mRNA was enriched using oligo-dT method and created remarkable 3’-bias due to the mRNA breakage as expected (Fig. 4F left of panel). Surprisingly, our protocol, which was developed for the non-degraded samples, could be directly used in the degraded samples without any change in experiment and analysis, and yield highly comparable gene expression values as non-degraded samples (Pearson R^2^>0.95, Fig. 4B). The rRNA depletion strategy avoids the 3’-bias (Fig. 4F middle of the panel), and also showed remarkable consistency to the non-degraded counterparts (R^2^>0.88, Fig. 4C), but still lower than the oligo-dT method.

**Figure 4:**
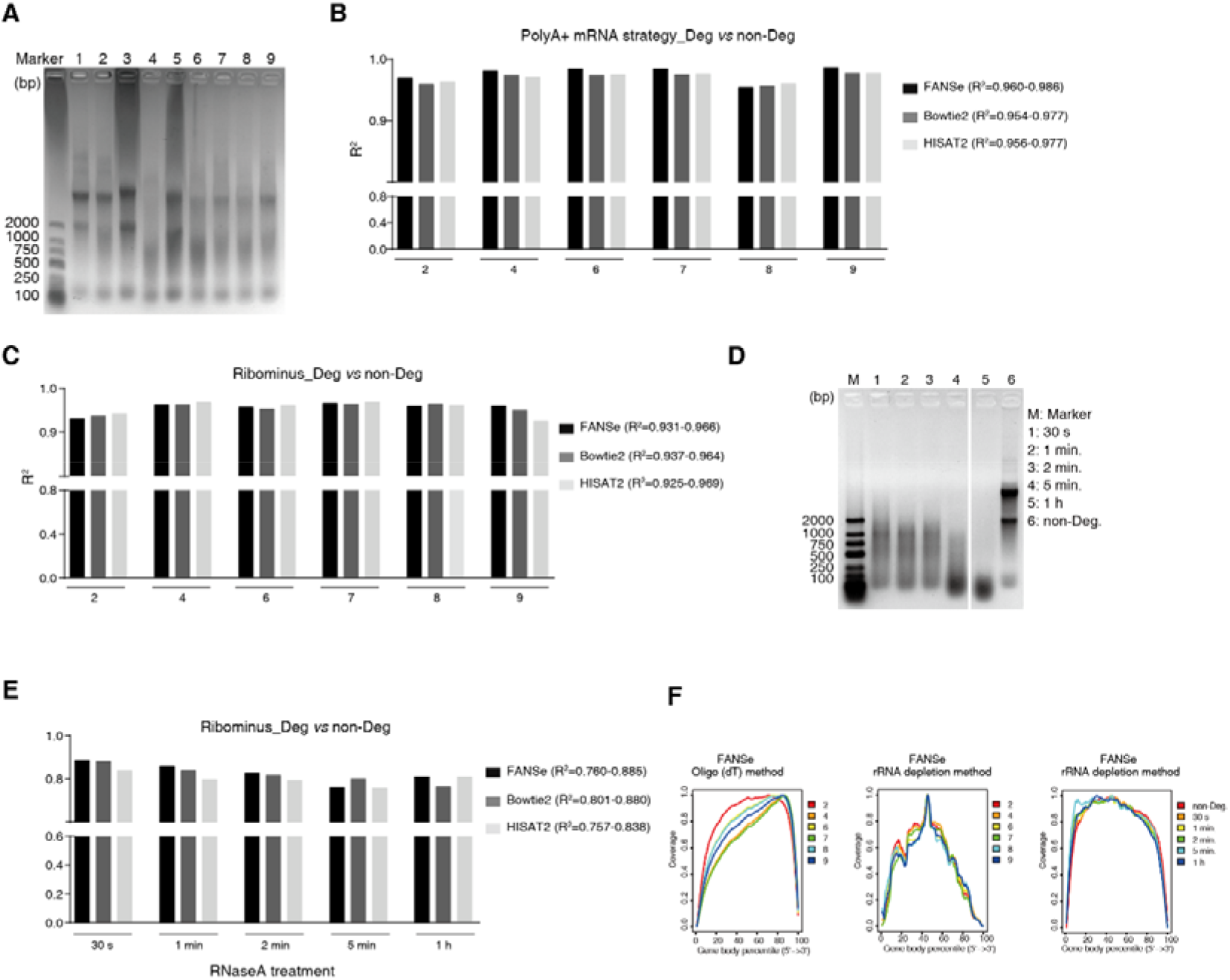
RNA-seq of degraded MHCC97H RNA samples. (A) The electrophoresis of the total RNA of MHCC97H, which was frozen and thawed for 10 cycles and exposed to air for prolonged time. (B) Correlation of the polyA+ transcriptome quantifications of the degraded and its corresponding non-degraded RNA samples. The raw sequencing datasets were analyzed using FANSe3, Bowtie2 and HISAT2, respectively. (C) Similar to Figure 4B, but using the Ribominus strategy. (D) Artificially degraded MHCC97H total RNA by adding RNase A for 30 seconds to 1 hour. (E) Similar to Figure 4C, for the RNaseA-degraded RNA. (F) Read coverage distribution along the mRNAs of the RNaseA-degraded RNA.

Most environmental RNases are exonucleases, which may remain the 3’-end of the mRNAs. However, endonucleases may degrade mRNA into smaller fragments. We added RNase A into the non-degraded MHCC97H total RNA and incubated for 30 seconds to 1 hour to create a series of endonuclease-degraded samples (Fig. 4D). Due to the endonuclease cleavage, sequencing library construction using oligo-dT strategy was failed (data not shown) but succeeded using rRNA depletion strategy. The coverage of the reads was much more uniform than the environmentally-degraded samples (Fig. 4F right of panel). However, the endonuclease-degraded samples showed considerably lower consistency compared to the non-degraded counterparts (Fig. 4E). Using FANSe3 algorithm, R^2^=0.760∼0.885; while using Bowtie2/HISAT2 + featureCounts, the correlation is even lower (R^2^=0.757∼0.880).

### The reproducible proteomics

The stable transcriptome of MHCC97H indicated also a stable proteome. However, translational regulation is the most significant regulatory level (*22*). Therefore, we first tested the stability of the MHCC97H translatome over subculture generations. The RNC-seq of the MHCC97H showed also very high mutual consistency (Pearson R^2^=0.948∼0.991, Fig. 5A). This made us to hypothesize that the proteome is also consistent over the generations. Indeed, using our protocol, the protein abundance detected using MS were comparable both in DDA mode (R^2^=0.934∼0.976) and in DIA mode (R^2^=0.942∼0.986), respectively (Fig. 5B). To test the variability contributed by the experimental procedures, we started from the same trypsin-digested sample and made 3 independent MS measurements (including LC-MS and data analysis). Such technical replicates yielded R^2^=0.945 and R^2^=0.975 in DDA and DIA modes, respectively (Fig. 5C). These results indicated that the variance contributed by the biological nature and trypsin digestion could be neglected in the DDA mode, and merely distinguishable in the DIA mode.

**Figure 5:**
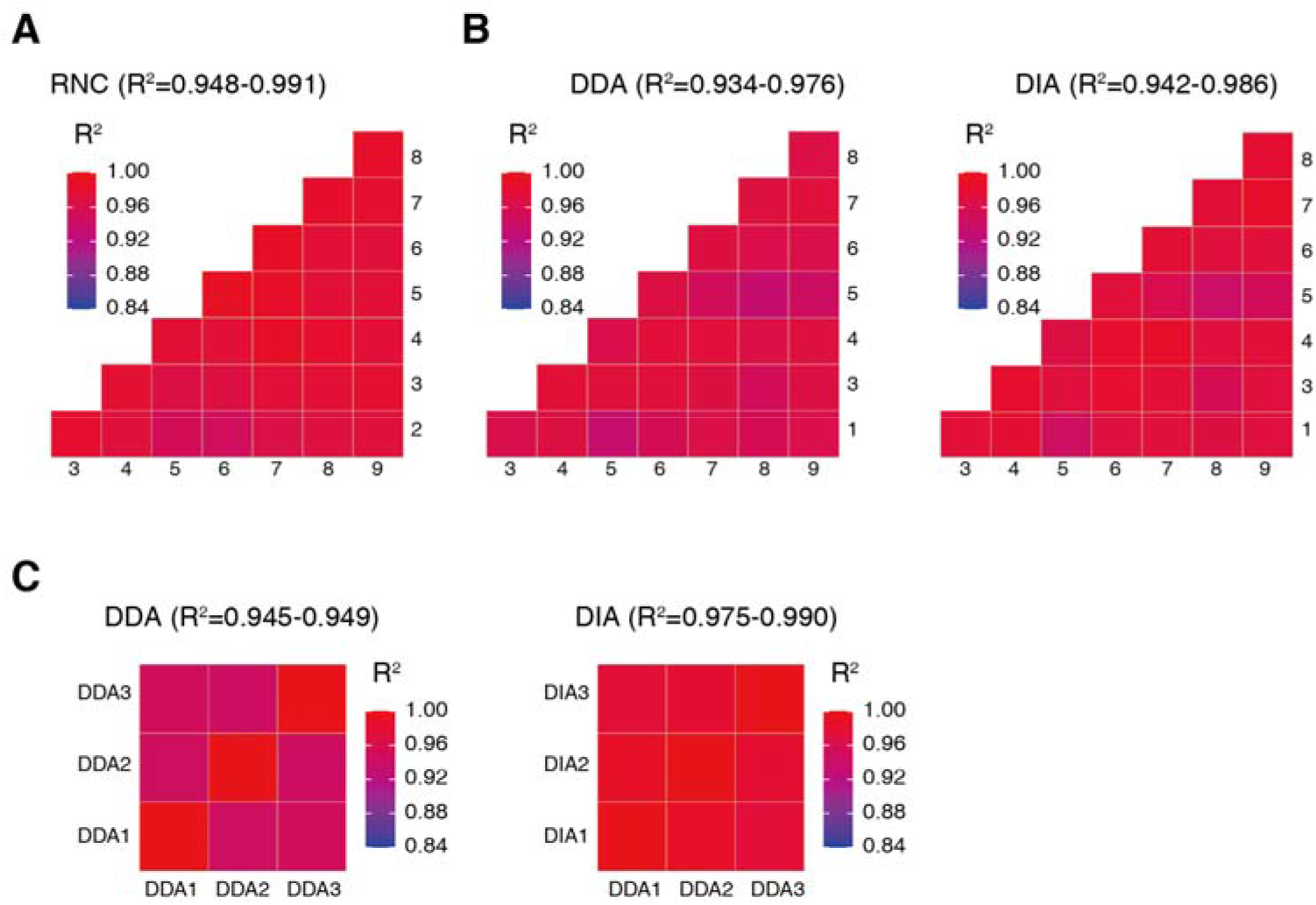
The stable translatome and proteome of MHCC97H during subculturing. (A) The RNC-seq of the MHCC97H of 7 subculturing generations. (B) The consistency of protein abundance of the MHCC97H detected by MS in DDA and DIA modes, respectively. (C) The technical triplicate of DDA and DIA modes of one MHCC97H total protein sample.

Next, we tested the robustness of the standard proteome sample across labs and instruments. We distributed the same batch of standard proteome sample to 4 labs (JNU, SCUT, DICP and BNU), which were over 2000 km away(Fig. 6A). The samples were shipped using ice boxes at 0 °C for 3 days. All labs followed the same protocol to process the samples. The only hardware differences were listed in Fig. 6B. The number of identified proteins were similar in the 4 labs (Supplementary Fig.2A∼B). The JNU lab yielded slightly more proteins due to the longer column, which provides higher chromatography resolution. The SCUT lab yielded fewer proteins due to the lower resolution and slower scanning speed of the mass spectrometer. However, the distribution of the isoelectric points of the identified protein showed no significant differences among these labs (Fig. 6C). The protein abundance measured by these labs were highly comparable (R^2^=0.925∼0.949, Fig. 6D left of panel). The variations of the DDA mode measurements were similar to the variations over the generations in one lab, demonstrating the high robustness of our SOP across the lab and instruments. In DIA mode, the labs with the same instruments showed highly similar results (R^2^=0.926), while the SCUT lab, which was equipped with another model of mass spectrometer showed remarkable lower consistency to the other two labs (R^2^=0.832) (Fig. 6D,right of panel), demonstrating that the instrument-specific bias cannot be neglected.

**Figure 6:**
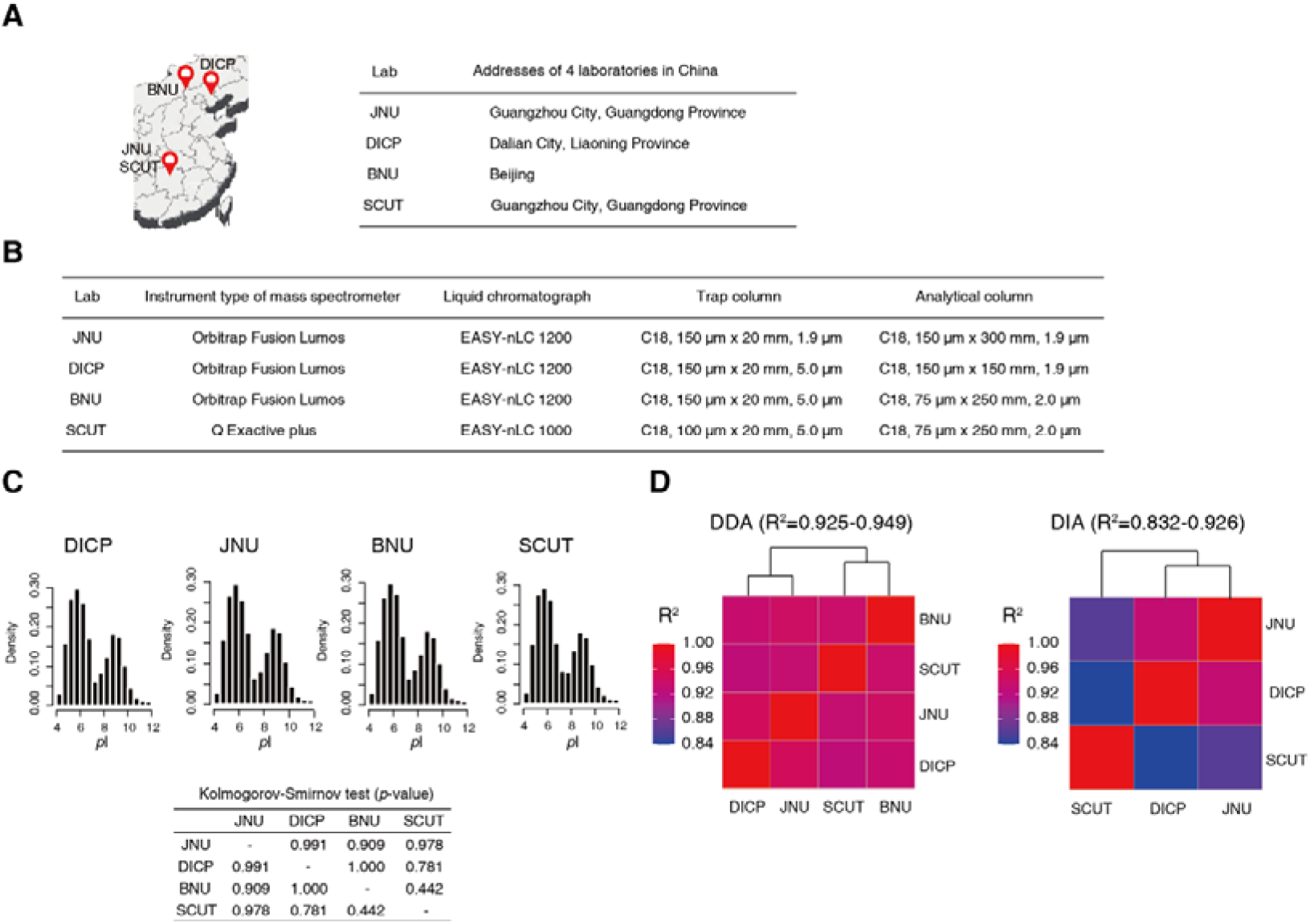
The repeatability of the MHCC97H proteome was verified in several laboratories. (A) The locations of the four laboratories in China (JNU, SCUT, DICP and BNU). (B) Hardware configuration of MS instruments in 4 labs (JNU, SCUT, DICP and BNU). (C) Distribution of isoelectric point of the same batch of MHCC97H protein samples identified by 4 laboratories, and their mutual Kolmogorov-Smirnov tests. (D) Mutual Pearson correlation of the protein abundances measured by 4 laboratories in DDA and DIA modes, respectively.

## Discussion

Ensuring reproducibility is an essential aspect of the omics era. A stable standard sample with omics-level complexity is necessary to assess and calibrate the entire RNA-seq and proteome MS workflows. We have shown that the human hepatocellular carcinoma cell line MHCC97H has a very stable transcriptome and proteome over 9 subculturing generations. In each subculturing generation, the cells can proliferate hundreds and thousands of times. Subculturing 9 generations can produce millions of tons of cells that allows constant industrial-scale production of such stable standard samples for many decades.

The purpose of the MHCC97H differs from the UHRR. The goal of the UHRR was to provide “hybridization signal at each microarray probe location (spot)” (*10*). To achieve this goal, the UHRR must cover the mRNAs of all coding genes, which will not be all expressed in any single tissue. Therefore, UHRR mixed 10 human cell lines which were originated from different tissues to cover as many mRNAs as possible. This goal does not require the stability in mRNA quantitation. Therefore, using the unstable cell lines such as HeLa is acceptable in UHRR, but not in the case of calibrating the workflow to yield high reproducibility.

An interesting finding was that the mild and natural RNA degradation by freeze-thaw cycles and environmental RNases did not affect the RNA quantification dramatically, even when using the identical polyA+ mRNA selection procedure that did not consider degradation at all (degraded vs. non-degraded R^2^=0.960∼0.986, Fig. 4B). The data of the degraded samples are directly comparable with the non-degraded ones. Such consistency was even better than using the Ribominus strategy (degraded vs. non-degraded R^2^=0.931∼0.971, Fig. 4C). This means that the former RIN (RNA integrity number)-based RNA quality criteria (*42, 43*) can be neglected in such cases. Despite of the obvious 3’-end bias of the reads from the degraded samples, the gene expression quantification (in RPKM) was not much affected, probably due to the universal and proportional loss of the 5’-end reads. This is a good news to the researchers who need to handle the degraded RNA samples due to the internal degradation, prolonged storage and transportation, fluctuated temperatures, etc. Also, the polyA+ mRNA selection should be a priority when possible. The artificial degradation by endonucleases caused lower correlation (degraded vs. non-degraded R^2^=0.760∼0.885, Fig. 4E), but still much higher than the correlation reported by other literatures (R^2^=0.37∼0.5) (*13*). These results showed the importance of a robust protocol.

The proteomics MS needs more steps than RNA-seq and thus were demonstrated quite difficult to reach high reproducibility (*4-6*). However, our results showed that the proteome quantification is highly reproducible in both DDA and DIA modes (average R^2^=0.967±0.016 for DDA and average R^2^=0.975±0.0114 for DIA, Fig. 5B). It has been thought that the DIA mode is more accurate and reproducible than the DDA (*44-46*). However, our results showed that the reproducibility of the DDA was similar to that of DIA in terms of biological replicates. The DDA mode generates slightly higher noise (R^2^=0.945∼0.949) than the DIA mode (R^2^=0.975∼0.990) in technical replicates (i.e. perform MS replicates using the same sample, Fig. 5C). However, such noise was covered by the biological noise (R^2^=0.934∼0.976, Fig. 5B left of panel). This also emphasized the importance of a standardized protocol.

In sum, we demonstrated that the MHCC97H cell line provides a stable transcriptome and proteome that serve as omics standards to assess and calibrate omics workflows. We also provided robust protocols to achieve good reproducibility. We believe that these efforts set a useful reference in the omics era.

## Methods

### Cell culture and materials

Hela, A549 cells were purchased from American Type Culture Collections (ATCC, Rockville, MD, USA) and authenticated by short tandem repeat profiling. the human hepatoma MHCC97H, HCCLM3, HCCLM6 cells were provided by Professor Yinkun Liu, Fudan University (*12*). All cells were detected free of mycoplasma during maintenance and upon experiments. These cells were cultured in DMEM medium with 10% FBS and 1% penicillin/streptomycin. All cell lines culture environment was 37 □, 5% CO2.

### RNA extraction

Cells from each generation were cultured to 80–90% confluency and washed twice with RNase-free PBS (LEAGENE, Beijing, China), and then isolated by using TRIzol RNA extraction reagent (Ambion, Austin,TX), following the manufacturer’s instructions.

### Ribosome-nascent chain complex-RNA (RNC-RNA) extraction

The RNC extraction was performed as we previously reported (PMID:23519614). In brief, cells from each generation were cultured to 80–90% confluency and were pre-treated with 100 mg/mL cycloheximide for 15 min, followed by pre-chilled PBS washes and addition of 2 mL cell lysis buffer (1% Triton X-100, 20 mM HEPES-KOH (pH 7.4), 15 mM MgCl_2_, 200 mM KCl, 100 mg/mL cycloheximide and 2 mM dithiothreitol (DTT)). After 30 min ice-bath, cell lysates were scraped and transferred to 1.5 mL RNase free tubes. Cell debris was removed by centrifuging at 16200 xg for 10 min at 4 °C. Supernatants were transferred on the surface of 20 mL of sucrose buffer (30% sucrose, 20 mM HEPES-KOH (pH 7.4), 15 mM MgCl_2_, 200 mM KCl, 100 mg/mL cycloheximide and 2 mM DTT). RNC was pelleted per ultra-centrifugation at 42500 xg for 5 h at 4 °C. Subsequently, RNC-RNA was extracted from RNC particles using TRIzol RNA extraction reagent following the manufacturer’s instructions.

### RNA degradation experiment

For RNA samples that are slightly degraded during RNA extraction, our study used two methods to construct the library, included Oligo (dT) method and rRNA depletion method. To evaluate the extent of degradation of each RNA sample, the GeneBody coverge.py of RNA-SeQC was used to evaluate the distribution of sequenced reads on the reference gene (*47, 48*).

The preparation method of RNA samples degraded by enzymes are as follows: First, we randomly selected a complete RNA sample without degradation and set it into six experimental groups (NC, 30 s, 1 min, 2 min, 5 min and 1 h), then took 2 ug RNA into each experimental group. Secondly, 1 ng RNase A was added to five experimental groups except the control group. The activity of RNase A was terminated by adding 0.5 U RNase OUT under the reaction time of 30 s, 1 min, 2 min, 5 min and 1 h, respectively. All experiments were operated at room temperature. The evaluation of the degradation was the same as described above.

### mRNA-seq and RNC-seq

For mRNA-seq, our study used two methods to construct the sequencing library, including Oligo (dT) method and the rRNA depletion method.

The sequencing libraries of Oligo (dT) were constructed using Library Preparation VATHS mRNA Capture Beads(Vazyme, N401-02, China)and MGIEasy RNA Library Prep Kit(MGI, A0210, China), following the manufacturer’s protocol. rRNA depletion sequencing libraries were also constructed using the MGIEasy RNA Library Prep Kit according to the manufacturer’s protocol too. Before the sequencing libraries were constructed, rRNA was removed from total RNA using RNaseH method as we previously reported (PMID: 31340039, PMID: 26926465).

For RNC-seq, only Oligo (dT) method was used for library construction, which was the same as mRNA-seq.

All the sequencing was performed on a MGISEQ-2000 sequencer (MGI, Shenzhen, China) for 210 cycles. The high-quality reads were subjected to the subsequent bioinformatics analysis. The adapter sequences were trimmed from the reads. Then reads were mapped to transcripts using the hyper-accurate mapping algorithm FANSe3 (*49, 50*) in the NGS analysis platform “Chi-Cloud” (http://www.chi-biotech.com). Gene expression levels were quantified using the rpkM method (*51*). Genes with at least 10 reads were considered quantifiable genes (*52*).

### Protein trypsin digestion

Cells of each generation were cultured to 80–90% confluency and treated with 0.25% trypsin-EDTA (Gibco, 25200056, USA), centrifuged at room temperature at 300 xg for 5 min, and washed with PBS (LEAGENE, Beijing, China) twice, and centrifuged to remove supernatant. Cells were dissolved in 1% SDS lysis buffer (Beyotime, P0013G, China) and the protein concentration was measured using a BCA kit (ThermoFisher, 23227, USA). The protein digestion was performed by filter-aided sample preparation (FASP) (*53*). In brief, protein samples were treated with 8M urea (8 M urea in 0.1 M Tris-HCl, pH 8.5) so that the final urea concentration is ≥4 M. Then add an appropriate amount of dithiothreitol solution (DTT) (Solarbio, D8220, China) to make its working concentration of 50 mM, incubate at 37 □ for 1 h. Then add an appropriate amount of iodoacetamide solution (IAA) (Merck, I6125, USA) to make its working concentration at 120-150 mM, and avoid light for 30 min at room temperature. All the solution was transferred to a 10KDa ultrafiltration tube (Merck, UFC501096, USA), centrifuged at 12000 xg and washed 3 times with 50 mM TEAB (ThermoFisher, 90114, USA). Trypsin (Promega, V5280, USA) was then added in a ratio of 1:40, and incubated overnight at 37 °C. The peptides were collected into low-binding collection tube (Thermo Scientific™, 88379, USA) by centrifugation at 12000 xg for 20 min. The filter tubes were washed twice with 50 mM TEAB by centrifugation at 12000 xg for 20 min. The flow-through peptides were collected, and then measured for the concentration using Pierce Quantitative Fluorometric Peptide Assay (Thermo Scientific™, 23290, USA). The peptides were then desalted using a Waters C18 columns (Waters, WAT054955, USA) following the steps below. First, the C18 columns were condition with 1 mL acetonitrile (ACN) (Thermo Scientific™, A955-4, USA), then equilibrated with 1 mL 5% ACN with 0.5% trifluoroacetic acid (TFA) (Macklin, T818782, China). The peptides which were acidized to pH 3-4 with TFA were loaded onto the C18 resin bed and then wash the C18 resin with 1mL 5% ACN with 0.5% TFA. Then the purified peptides were eluted with 70% ACN and collected into low-binding tubes, dried by vacuum centrifugation and then stored at -80 °C.

### Data-Dependent Acquisition (DDA) mass spectrometry

For DDA analysis, data were collected by Q Exactive mass spectrometer equipped with EASY-nLC 1000 system (Thermo Fisher Scientific, USA) and Orbitrap Fusion Lumos mass spectrometer equipped with EASY-nLC 1200 system (Thermo Fisher Scientific, USA) respectively.

#### QE parameter setting

2 µg of peptides containing 1 µL of standard peptides (iRT kit) (Biognosys, Ki-3002-2, Switzerland) were loaded on a nano trap column (Acclaim PepMap100 C18, 100 µm x 20 mm, 5 μm, Thermo Scientific™, AAA-164564, USA), and then separated onto an analytical column (PepMap C18, 75 μm x 250 mm, 2 μm, Thermo Scientific™, 164941, USA) using a 120 min linear gradient (solvent A: 98% H2O, 2% ACN, 0.1% FA; solvent B: 98% ACN, 2% H2O, 0.1% FA) at a flow rate of 300 nL/min. The detailed solvent gradient was as follows: 3–7% B, 4 min; 7– 18% B, 70 min; 18–25% B, 20 min; 25–35% B, 16 min; 35–40% B, 1 min;40–90% B, 9 min. The mass spectrometer was operated in data dependent top-20 mode with the following settings: MS1 scan was acquired from 400 to 1200 m/z with a resolution of 70000, the auto gain control (AGC) was set to 3e6, and the maximum injection time was set to 60 ms. MS2 scans were performed at a resolution of 17500 with an isolation window of 1.6 m/z and higher collision energy dissociation (HCD) at 32%, the AGC target was set to 5e5, and the maximal injection time was 50 ms, the loop count was set to 20 and the normalized collision energy (NCE) was 27%, with dynamic exclusion of 30 s.

#### Orbitrap Fusion Lumos parameter setting

2 µg of peptides containing 1 µL of standard peptides (iRT kit) (Biognosys, Ki-3002-2, Switzerland) were loaded on a nano trap column (C18, 150 µm x 20 mm, 1.9 μm, homemade), and then separated onto an analytical column (C18, 150 μm x 300 mm, 1.9 μm, homemade) using a 120 min linear gradient (solvent A: 98% H2O, 2% ACN, 0.1% FA; solvent B: 98% ACN, 2% H2O, 0.1% FA) at a flow rate of 600 nL/min. The detailed solvent gradient was as follows: 5–12% B, 28 min; 12–24% B, 58 min; 24–38% B, 25 min; 38–95% B, 1 min; 95% B, 8 min. The data dependent mass spectrometer acquisition was operated in top-speed mode with the following settings: MS1 scan was acquired from 350 to 1500 m/z with a resolution of 120k, the auto gain control (AGC) was set to 4e5, and the maximum injection time was set to 50 ms. MS2 scans were performed at a resolution of 15k with an isolation window of 1.6 m/z, the higher collision energy dissociation (HCD) collision energy was set to 31%, the AGC target was set to 5e4, and the maximal injection time was set to 50 ms, the cycle time was set to 3s with a dynamic exclusion of 30 s.

### Data-Independent Acquisition mass spectrometry

DIA data were also collected using QE and Orbitrap Fusion Lumos respectively.

#### QE parameter setting

2 µg of peptides containing 1 µL of standard peptides of each sample were analyzed in DIA method. The liquid conditions were the same as those of the DDA model. For MS acquisition, the MS1 resolution was 70000, and the MS2 resolution was set to 17500. The m/z range covered from 400 to 1200 m/z and was separated into 30 variable acquisition windows. The full scan AGC target was set to 3e6, with an injection time of 60 ms. DIA settings included NCE of 27%, AGC target of 1e6, auto maximum injection time.

#### Orbitrap Fusion Lumos parameter setting

2 µg of peptides containing 1 µL of standard peptides of each sample were analyzed in DIA method. The liquid chromatography conditions were the same as those of the DDA model. For MS acquisition, the MS1 scan at resolution 120000 with an AGC target of 4e5 and max injection time of 50 ms in the mass range of 350 to 1250 m/z, followed by 40 DIA scans with segment widths adjusted to the precursor density. The MS2 scan resolution was set to 30k with an AGC target of 5e5 and max injection time of 50 ms. The HCD collision energy was set to 31%.

### Database search

MaxQuant (version 1.5.7.4) was used for DDA data search. The common search parameters: type, standard, multiplicity 1; digestion, digestion mode(specific), enzyme, trypsin/P; variable modification, oxidation(M), acetyl (protein N-term), max number of modifications per peptide 5, two missed cleavage sites were allowed; label-free quantification, LFQ, LFQ min ratio count 2, fast LFQ, LFQ min number of neighbors 3, LFQ average number of neighbors 6; instrument, orbitrap; fixed modification, carbamidomethyl (C); two missed cleavage sites were allowed. Adv. identification, second peptides; protein quantification, label min ratio count 2, peptides for quantification unique+ razor, discard unmodified counterpart peptides, advanced ratio estimation; label free quantification, stabilize large LFQ ratios, require MS/MS for LFQ comparisons, iBAQ; Specific to MS analyzers, mass tolerance parameters were set as follows: MS/MS-TMS,TMS MS/MS match tolerance 0.5 Da, TMS MS/MS de novo tolerance 0.25 Da, TMS MS/MS deisotoping tolerance 0.15 Da, TMS top peaks per Da interval 8, TMS top x mass window 100 Da; MS/MS-TOF,TOF MS/MS match tolerance 40 ppm, TOF MS/MS de novo tolerance 0.02 Da, TOF MS/MS deisotoping tolerance 10 Da, TOF top peaks per Da interval 10, TOF top x mass window 100; MS/MS-Unknow, unknown MS/MS match tolerance 0.5 Da, unknown MS/MS de novo tolerance 0.25 Da, unknown MS/MS deisotoping tolerance 0.15 Da, unknown MS/MS top peaks per Da interval 8, unknown MS/MS top x mass window 100 Da. We adopted the criteria for confident identification with false discovery rate (FDR)<0.01, at peptide and protein levels. Data from label-free MS analyses were searched with all three search engines. DIA data search using the direct DIA module of Spectronaut powered by Pulsar 14.2.200619.47784 (Biognosys, Switzerland). The common search parameters: the search allowed 2 missed cleavage, and the enzyme was set to trypsin/P. Carbamidomethyl (cysteine) was allowed as a fixed modification, and Oxidation (methionine), Acetyl (Protein N-term) were allowed as variable modifications. The mass tolerance for matching precursor and fragment ions: dynamic; The identification was performed using 0.01 FDR threshold at peptide, protein and PSM levels. The calibration was set to non-linear iRT calibration with precision iRT enabled. The identification was performed using 0.01 Q-value (adjust P value) cutoff on precursor and protein level while the maximum number of decoys was set to a fraction of 0.75 of library size. For quantification interference correction was enabled with at least three fragment ions used per peptide, major and minor group quantities were set to mean peptide and mean precursor quantity, respectively with Top3 group selection each. Quantity was determined on MS2 level using the area of XIC peaks with enabled cross run normalization.

The Database of Uniprot-Human-Filtered-Reviewed-Yes -UP000005640_9606.fasta was used for all database search.

### RNA and protein quantification

For mRNA-seq and full-length translating mRNA-seq (RNC-seq) data, our study used three different algorithms, including FANSe3, HISAT2 and Bowtie2. The sequence mapping of FANSe3 can be referenced to human transcriptome database, while the sequence mapping of HISAT2 and Bowtie2 can be referenced to human genome database. The mRNA abundance was normalized using both rpkM (reads per kilobase per million reads).

For protein quantification analysis, label-free MS data were quantified with the iBAQ (intensity-based absolute quantification) algorithm as provided in MaxQuant. The missing values of protein quantitative data were deleted and after the median normalization.

## Supporting information

Successful extraction of complete RNA from 5 cell lines

Number of proteins in different laboratories in DDA and DIA modes, respectively

RNA-seq quality contorl for all cell lines

Details of transcriptome data for 5 cell lines

Details of transcriptome data for MHCC97H stability assessment

Details of MHCC97H transcriptome data using different library construction methods

Details of MHCC97H translatomics data and proteomic data

Details of MHCC97H proteomic data in several laboratories

## Data availability

All the sequencing data sets are available at NGDC-GSA for Human (Submission of HRA: HRA001521, HRA001871). All the MS raw data are publicly available in NGDC-OMIX (Submissions of OMIX: OMIX882, OMIX883, OMIX002283). Details of all omics data are shown in Supplementary table s2-Supplementary table s6.

## Code availability

No code is used in this manuscript.

## Acknowledgements

We would like to thank Chi-biotech Co. Ltd and SAGENE Co. Ltd for their contributions to the multi-platform and multi-center data generation of transcriptome.

## Author contributions

G.Z., S.H.L. and Y.C. conceived the project, designed the experiments and co-supervised the study, S.H.L. and Y.C. performed cell culture, S.H.L. performed mass spectrometry experiments with assistance by Z.H.S., H.L. and T.K.Z. performed the mRNA-seq experiments with assistance by Y.C. and J.J.J, H.L. performed the analysis of all omics data with assistance by S.H.L., Y.C. performed the RNC-seq experiments, H.M.Y., H.L.D., Y.H.G., Y.T.L., X.Z.P., W.L.Z. and S.Y.F. jointly performed the multi-center and multi-platform mass spectrometry experiments, Q.Y.H. providing mass spectrometry resource, G.Z. and S.H.L. wrote the manuscript. All authors read and approved the final manuscript.

