## Supplementary figures and images for "A stable reference human transcriptome and proteome as a standard for reproducible omics experiments"

### Number of proteins in different laboratories in DDA and DIA modes, respectively

#### Supplementary Figure 2

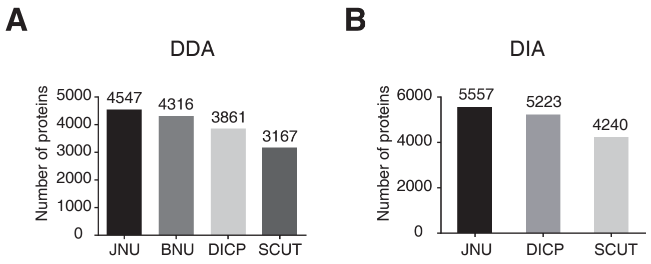

### Successful extraction of complete RNA from 5 cell lines

#### Supplementary Figure 1

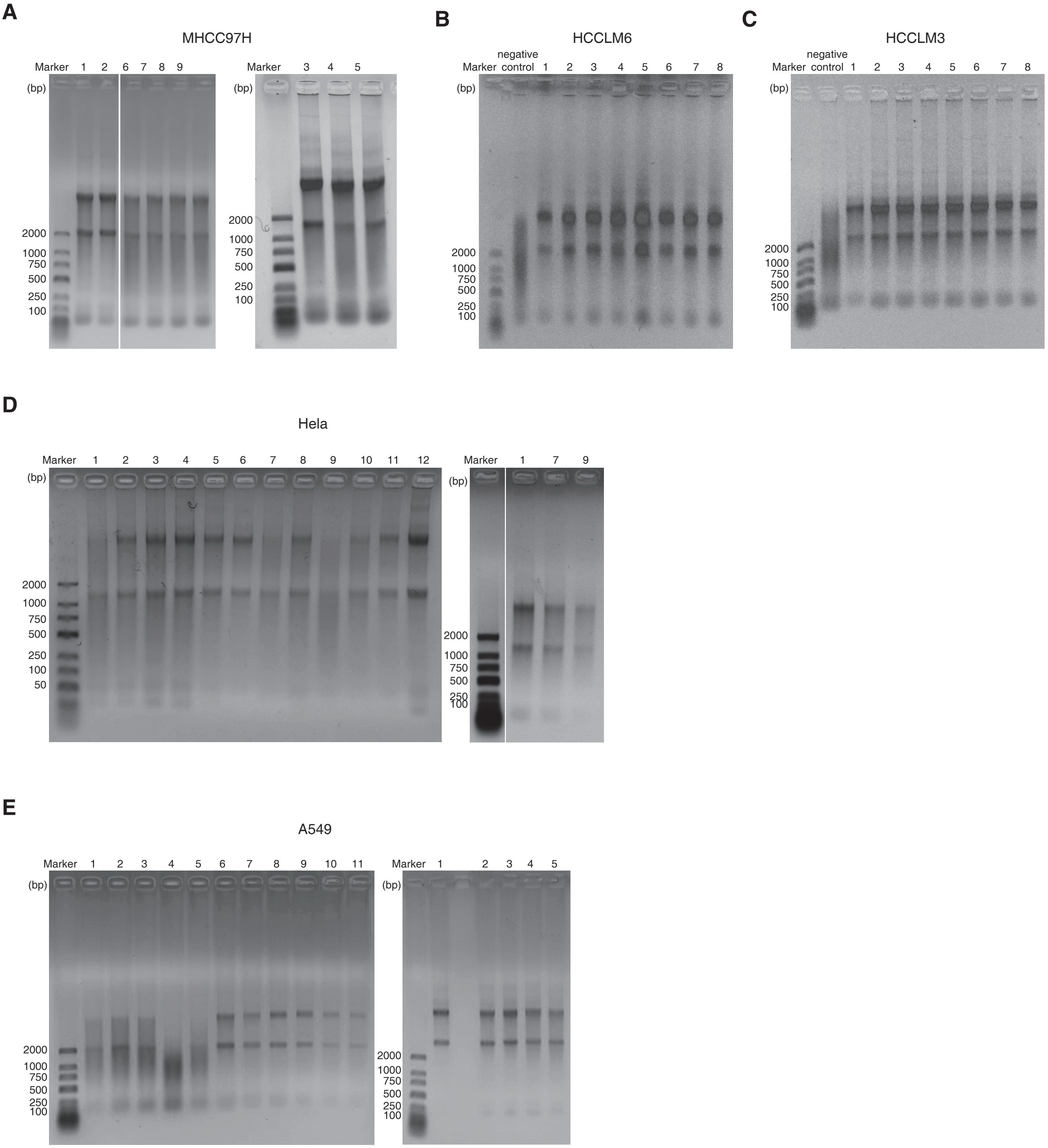
