## Supplementary material for "A stable reference human transcriptome and proteome as a standard for reproducible omics experiments": RNA-seq quality contorl for all cell lines

| Supplementary table s1: RNA-seq quality contorl for all cell lines. | | | | | |
|---|---|---|---|---|---|
| cell line | index | generation | Q30 (%) | rRNA reads ratio | mapping ratio |
| MHCC97H | 81 | g-1 | 88.18 | 1.26% | 78.63% |
| | 82 | g-2 | 88.53 | 0.67% | 70.83% |
| | 83 | g-3 | 89.23 | 0.67% | 68.98% |
| | 84 | g-4 | 88.13 | 2.43% | 80.84% |
| | 85 | g-5 | 88.62 | 2.00% | 78.14% |
| | 86 | g-6 | 89.02 | 2.56% | 79.19% |
| | 87 | g-7 | 88.56 | 2.01% | 80.74% |
| | 88 | g-8 | 89.13 | 1.85% | 74.03% |
| | 73 | g-9 | 88.90 | 2.07% | 79.44% |
| A549 | 89 | g-1 | 91.21 | 3.52% | 79.14% |
| | 90 | g-2 | 88.48 | 2.98% | 81.77% |
| | 91 | g-3 | 85.61 | 2.66% | 82.31% |
| | 92 | g-4 | 87.55 | 2.73% | 81.90% |
| | 93 | g-5 | 89.96 | 3.24% | 81.91% |
| | 94 | g-6 | 80.83 | 3.61% | 77.97% |
| | 95 | g-7 | 83.92 | 4.33% | 78.96% |
| | 96 | g-8 | 82.88 | 3.97% | 78.31% |
| | 85 | g-9 | 86.11 | 3.37% | 78.45% |
| | 86 | g-10 | 89.68 | 3.98% | 80.90% |
| | 87 | g-11 | 83.68 | 3.12% | 82.73% |
| Hela | 73 | g-1 | 87.70 | 6.89% | 81.50% |
| | 74 | g-2 | 90.64 | 7.03% | 82.28% |
| | 75 | g-3 | 88.13 | 3.31% | 82.20% |
| | 76 | g-4 | 86.87 | 3.99% | 82.17% |
| | 77 | g-5 | 84.97 | 4.56% | 81.83% |
| | 78 | g-6 | 90.45 | 5.34% | 79.49% |
| | 79 | g-7 | 85.35 | 3.99% | 80.87% |
| | 80 | g-8 | 89.54 | 3.45% | 78.79% |
| | 81 | g-9 | 81.60 | 5.19% | 82.66% |
| | 82 | g-10 | 87.41 | 3.99% | 80.47% |
| | 83 | g-11 | 88.14 | 4.58% | 82.03% |
| | 84 | g-12 | 82.46 | 4.05% | 76.56% |
| HCCLM3 | 66 | g-1 | 94.70 | 8.64% | 82.68% |
| | 67 | g-2 | 94.53 | 11.60% | 78.03% |
| | 68 | g-3 | 94.44 | 8.17% | 80.37% |
| | 69 | g-4 | 94.81 | 1.88% | 83.31% |
| | 70 | g-5 | 95.07 | 7.06% | 80.01% |
| | 71 | g-6 | 94.66 | 6.54% | 80.01% |
| | 72 | g-7 | 94.77 | 6.71% | 80.27% |
| | 75 | g-8 | 95.55 | 21.35% | 82.41% |
| HCCLM6 | 82 | g-1 | 94.56 | 25.29% | 86.93% |
| | 83 | g-2 | 95.59 | 26.95% | 77.97% |
| | 84 | g-3 | 95.54 | 33.65% | 86.04% |
| | 85 | g-4 | 95.77 | 17.51% | 82.01% |
| | 86 | g-5 | 95.76 | 17.95% | 79.76% |
| | 87 | g-6 | 95.51 | 5.50% | 81.69% |
| | 88 | g-7 | 95.57 | 6.41% | 84.03% |
| | 76 | g-8 | 95.63 | 16.90% | 77.97% |
